# Coordinated regulation of Mdr1- and Cdr1-mediated protection from antifungals by the Mrr1 transcription factor in emerging *Candida* spp

**DOI:** 10.1101/2025.05.04.652153

**Authors:** Dhanabala-Subhiksha Rajesh-Khanna, Carolina G. Piña Páez, Elora G. Dolan, Kiran S. Mirpuri, Jason E. Staijch, Deborah A. Hogan

## Abstract

Infections caused by the emerging pathogenic yeast *Clavispora (Candida) lusitaniae* can be difficult to manage due to multi-drug resistance. Resistance to the frontline antifungal fluconazole (FLZ) in *Candida* spp. is commonly acquired through gain-of-function (GOF) mutations in the gene encoding the transcription factor Mrr1. These activated Mrr1 variants enhance FLZ efflux via upregulation of the multi-drug transporter gene *MDR1*. Recently, it was reported that, unlike in the well-studied *Candida albicans* species, *C. lusitaniae* and *Candida parapsilosis* with activated Mrr1 also have high expression of *CDR1*, which encodes another multi-drug transporter with overlapping but distinct transported substrate profiles and Cdr1-dependent FLZ resistance. To better understand the mechanisms of Mrr1 regulation of *MDR1* and *CDR1*, and other co-regulated genes, we performed CUT&RUN analysis of Mrr1 binding sites. Mrr1 bound the promoter regions of *MDR1* and *CDR1* as well as *FLU1*, which encodes another transporter capable of FLZ efflux. Mdr1 and Cdr1 independently contributed to the decreased susceptibility of the *MRR1^GOF^* strains against diverse clinical azoles and other antifungals, including 5-flucytosine. A consensus motif, CGGAGWTAR, enriched in Mrr1-bound *C. lusitaniae* DNA was also conserved upstream of *MDR1* and *CDR1* across species including *C. albicans*. CUT&RUN and RNA-seq data were used to define the Mrr1 regulon which includes genes involved in transport, stress responses, and metabolism. Activated and inducible Mrr1 bound similar regions in the promoters of Mrr1 regulon genes. Our studies provide new evolutionary insights into the coordinated regulation of multi-drug transporters and potential mechanism(s) that aid secondary resistance acquisition in emerging *Candida*.

**SIGNIFICANCE:** Understanding antifungal resistance in emerging *Candida* pathogens is essential to manage treatment failures and guide the development of new therapeutic strategies. Like other *Candida* species, the environmental opportunistic fungal pathogen *Clavispora* (*Candida*) *lusitaniae* can acquire resistance to the antifungal fluconazole by overexpression of the multi-drug efflux pump Mdr1 through gain-of-function mutations in the gene encoding the transcription factor Mrr1. Here, we show that *C. lusitaniae* Mrr1 also directly regulates *CDR1,* another major multi-drug transporter gene, along with *MDR1.* In strains with activated Mrr1, upregulation of *MDR1* and *CDR1* protects against diverse antifungals potentially aiding the rise of other resistance mutations. Mrr1 also regulates several stress response and metabolism genes thereby providing new perspectives into the physiology of drug-resistant strains. The identification of an Mrr1 binding motif that is conserved across strains and species will advance future efforts to understand multi-drug resistance across *Candida* species.

## INTRODUCTION

Invasive or systemic candidiasis affects over 1.5 million people each year with high rates of mortality (1) and localized *Candida* infections have high economic and quality of life burdens. While *Candida albicans* is the major causative agent of *Candida* infections, other non-albicans *Candida* such as *Clavispora* (*Candida*) *lusitaniae* are garnering attention for increased incidence and drug susceptibility profiles (2). *C. lusitaniae* can establish difficult-to-treat infections in immunocompromised individuals (3–9). Unlike *Candida* spp. that are found largely within the human microbiome, *C. lusitaniae* appears to have a flexible physiology that allows it to occupy environmental, agricultural and human-associated niches (10). Much like its phylogenetic neighbor *Candidozyma* (*Candida*) *auris*, *C. lusitaniae* also can exhibit resistance to any of the three major antifungal classes – polyenes, echinocandins, and azoles such as fluconazole (FLZ) – within days of treatment (9, 11–15).

*C. lusitaniae,* like other *Candida* spp., gains resistance to FLZ through several mechanisms (12) including mutation of the FLZ target Erg11 or the acquisition of gain-of-function (GOF) mutations in the gene encoding for the multi-drug resistance regulator Mrr1. Mrr1^GOF^ variants have constitutive activity and upregulate the expression of the multi-drug efflux pump gene *MDR1*. The major facilitator superfamily (MFS) transporter Mdr1 is conserved across *Candida* species and has promiscuity for structurally and functionally distinct substrates including FLZ, bacterial phenazines and salivary antimicrobial peptides like histatins (16–18). Mrr1-dependent transcription of *MDR1* and other genes can be induced by xenobiotics such as benomyl (19–23) and the metabolite methylglyoxal (24). Across *Candida* species, Mrr1^GOF^ variants also co-regulate the expression of *MDR1* and putative methylglyoxal dehydrogenases (23, 25–28) and confer fitness advantages outside of FLZ resistance in *C. lusitaniae* (24).

In Demers *et al.* (21), we described the repeated selection for *MRR1^GOF^* mutations in clinical isolates recovered from a chronic lung infection of *C. lusitaniae*. These mutations evolved in a FLZ-naïve environment suggesting there are unrecognized roles for *MRR1* in host adaptation. Genomic analyses of these isolates identified secondary suppressor mutations that either attenuated constitutive activity or restored the inducible Mrr1 phenotype. The regain of inducibility underscores that there are benefits associated with an inducible Mrr1 phenotype as well. The physiology of strains with an inducible and activated Mrr1 seems to be vastly different as over 90 targets including those reported in other species such as *MDR1* and methylglyoxal dehydrogenase-encoding *MGD1* and *MGD2* as well as novel unanticipated targets like *CDR1*, encoding for a multi-drug transporter, were differentially expressed in a transcriptomic analysis of the different *MRR1* alleles (18, 21).

The ATP-binding cassette (ABC) superfamily transporter Cdr1 is a well-studied *Candida* multi-drug efflux pump that is conserved across species and exports a wide range of substrates including Mdr1 targets like FLZ as well as distinct ones like rhodamine-6-G (29–31). In *C. albicans*, *CDR1* expression is regulated by the zinc-cluster transcription factor Tac1 and GOF mutations in the *TAC1* gene is another mechanism of FLZ resistance (27, 32, 33). However, recent studies in emerging *Candida* spp. Including *C. lusitaniae* and *C. parapsilosis* have shown Mrr1-dependent changes in *CDR1* expression (34, 35). Here, we address this altered regulation of *CDR1* and its effect on strains with constitutively active Mrr1 in *C. lusitaniae*.

In this study, we report that *C. lusitaniae* Mrr1 directly regulates both *MDR1* and *CDR1* and that this coordinate regulation of Mdr1 and Cdr1 contributes to decreased sensitivity to multiple clinical and environmental antifungals. Further, analysis of Mrr1-DNA interactions found that Mrr1 directly regulates genes involved in cellular processes beyond drug transport by binding to a consensus Mrr1 motif that is conserved in different species. We also demonstrate that Mrr1 activation state does not alter its DNA localization at these targets. While this model for *MDR1* and *CDR1* regulation differs from that which has been described in *C. albicans*, the findings in *C. lusitaniae* are consistent with recent reports in diverse *Candida* species including *C. auris*. Our findings suggest that the rise of drug resistant lineages may be aided by the coordinated regulation of two drug resistance factors under Mrr1 and that plasticity in drug resistance regulation could be instrumental in the development of multi-drug resistant species.

## RESULTS

### *C. lusitaniae* Mrr1 effects on expression of multiple transporters involved in drug resistance

We previously characterized the *C. lusitaniae* clinical isolate strain U04 and its *mrr1*Δ derivative complemented with either *MRR1^ancestral^*, which confers Mrr1 activity typical of most *C. lusitaniae* isolates, or *MRR1^Y813C^*, which confers constitutive Mrr1 activity that renders cells resistant to FLZ (21). Published transcriptomic comparisons of these strains revealed significantly higher levels of *MDR1* (*CLUG_01938_39* (18)), *CDR1* (*CLUG_03113* (34)) and *CLUG_05825* (*FLU1*), all of which encode drug efflux proteins, when Mrr1 was constitutively active (21) (Fig 1A). Here on, we refer to *CLUG_05825* as *FLU1*, as the protein sequence encoded by this gene had the highest similarity by BLAST to the homolog encoded by *C7_01520W_A* of *C. albicans* SC5314.

**Figure 1:**
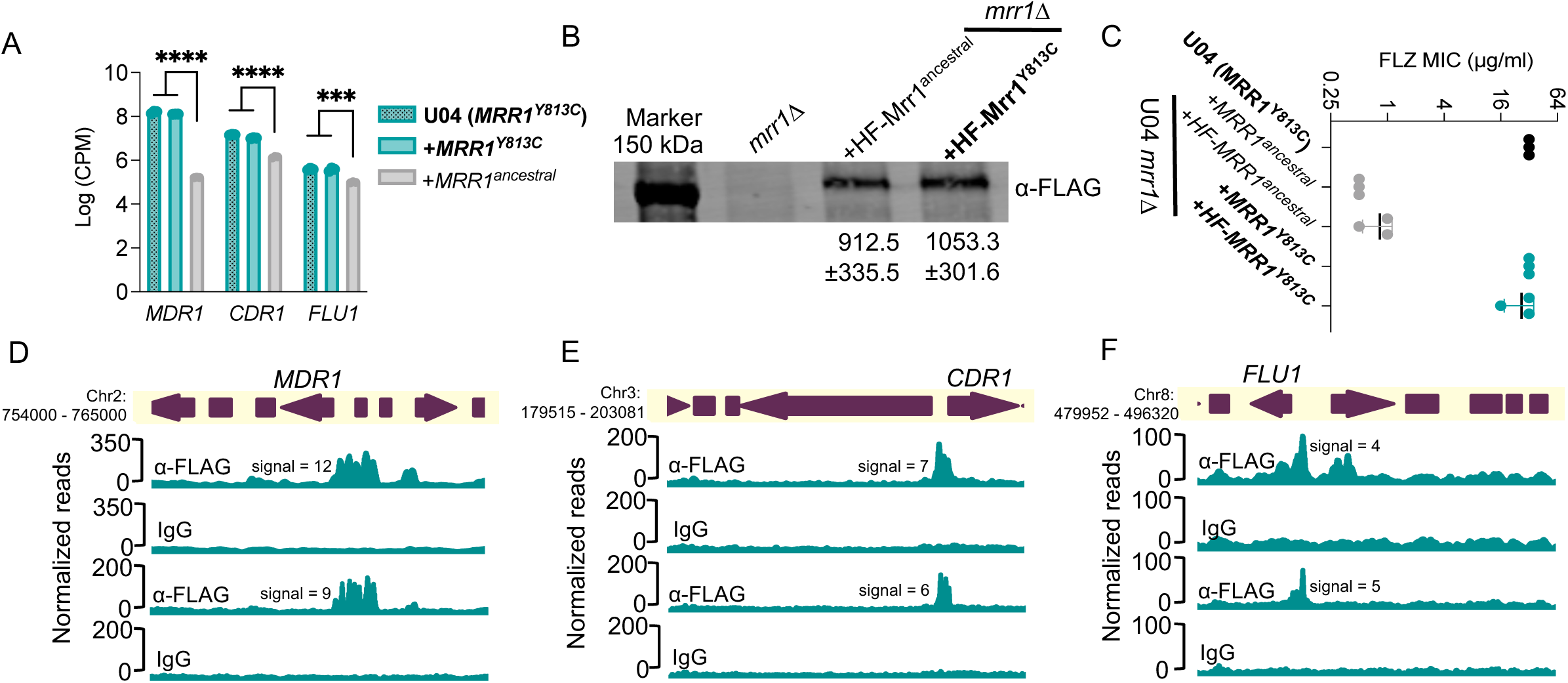
Biochemical and phenotypic analysis of HF-tagged Mrr1 and binding profiles of constitutively active Mrr1. (A) Log counts per million (CPM) values of *MDR1 (CLUG_01938_39)*, *CDR1 (CLUG_03113)* and *FLU1 (CLUG_05825)* from RNA-seq analysis of Demers *et al.* (21) comparing U04 clinical isolate (native allele *MRR1^Y813C^*) and U04 *mrr1*Δ complemented with either *MRR1^Y813C^* or *MRR1^ancestral^*. Ordinary one-way ANOVA and Dunnett’s multiple comparisons testing with a single pooled variance were used to evaluate the statistical significance for each gene. ***, p<0.001; ****, p<0.0001. (B) Western blot of whole cell protein lysates of U04 strains expressing N-terminal 6xHis-3xFLAG-tagged Mrr1 (HF-Mrr1) variants. HF-Mrr1 was probed using an α-FLAG antibody. Mean ± SD of HF-Mrr1 band intensities normalized to total protein (n= 4 biological replicates). (C) FLZ MIC of U04 clinical isolate and U04 *mrr1*Δ complemented with untagged or *HF-MRR1* was determined by broth microdilution assays. The data represent the mean ± SD from three independent experiments. There were no significant differences observed between data from strains with untagged Mrr1 variants and data from strains with their respective HF-tagged counterparts. (A – C) Strains with constitutive Mrr1 activity are in bold. (D-F) HF-Mrr1^Y813C^ CUT&RUN read coverage plots normalized per 20 bp bin size. Chromosomal positions of regions containing *MDR1, CDR1* and *FLU1* and adjacent genes are represented to scale with boxes and arrows. Peaks from HF-Mrr1^Y813C^-bound DNA recovered by an α-FLAG antibody and for the non-specific binding control recovered via IgG are shown. Signal indicates the average read density in α-FLAG relative to IgG within the peak region. Two independent experiments were performed and both are shown.

### Construction and activity of epitope-tagged Mrr1 variants

To investigate if *C. lusitaniae* Mrr1 regulation of these transporters was direct, we analyzed Mrr1-DNA interactions. We first generated N-terminal 6xHis-3xFLAG (HF)-tagged versions of different Mrr1 variants. HF-Mrr1-encoding alleles were expressed from the native *MRR1* promoter after introduction into the U04 *mrr1*Δ mutant background. We found that HF-Mrr1^ancestral^ and HF-Mrr1^Y813C^ were stably produced, and both were detected at a slightly higher molecular weight (150 kDa) than the predicted ∼140 kDa. This band was absent in the western blot of samples from the U04 *mrr1*Δ strain (Fig. 1B). We did not observe any significant differences in Mrr1 levels between strains expressing HF-Mrr1^ancestral^ and HF-Mrr1^Y813C^ (Fig. 1B). We compared the activities of the HF-Mrr1 variants to their untagged counterparts by evaluating the minimum inhibitory concentration (MIC) of FLZ in strains with either HF-tagged or untagged Mrr1 variants (Fig. 1C). The U04 *mrr1*Δ strain with untagged *MRR1^Y813C^* had a MIC that was 64-fold higher than the strain with untagged *MRR1^ancestral^* (FLZ MIC of 32 µg/ml vs 0.5 µg/ml) (21). The U04 *mrr1*Δ strain complemented with *HF-MRR1^Y813C^* had a similarly high MIC relative to the strain with *HF-MRR1^ancestral^* (Fig. 1C). Thus, the N-terminal HF-tag did not affect Mrr1 function.

### Analysis of Mrr1^Y813C^-DNA localization in *C. lusitaniae*

We evaluated genome-wide binding of HF-Mrr1^Y813C^ using Cleavage Under Targets and Release Using Nuclease (CUT&RUN) in two independent experiments (36). An α-FLAG antibody (Ab) was used for the enrichment of HF-Mrr1 bound DNA, and an IgG Ab was used to assess non-specific binding. The recovered DNA was sequenced and aligned to the genome of *C. lusitaniae* strain L17 (NCBI accession: ASM367555v2). Both U04 and L17 were isolated from the same clinical sample and differ by only ∼108 Single Nucleotide Polymorphisms and ∼130 Insertions/Deletions (18). The genome of L17 was utilized as it is a highly accurate genome produced by sequencing and assembly of reads obtained using Oxford Nanopore long-read and Illumina technologies. Genomic regions that showed significant fold enrichment in DNA recovered from the α-FLAG when compared to the IgG control of each sample are represented as peaks and indicate HF-Mrr1^Y813C^ interaction sites (Fig. 1D). The average enrichment of reads in α-FLAG relative to the IgG background within an identified peak region was quantified as peak signal (37). Peaks were filtered using a peak signal cutoff of 2, a false discovery rate (FDR) of <0.05 and a genomic position within 1 kb of an open reading frame (ORF). Approximately 329 CUT&RUN peaks were identified (File S1).

The upstream regions of *MDR1*, *CDR1* and *FLU1* all showed strong evidence for Mrr1^Y813C^ binding. The upstream region of the *MDR1* ORF containing its promoter had a significant HF-Mrr1^Y813C^ peak with an average signal of 10 (Fig. 1D). The HF-Mrr1^Y813C^ peak associated with the *MDR1* ORF spanned ∼1.5 kb and extended into the neighboring coding regions of *MDR1* (Fig. 1D). Thus, as in *C. albicans* (20, 38), *C. lusitaniae* Mrr1 bound directly upstream of *MDR1*. The regions upstream of *CDR1* also had a significantly enriched CUT&RUN peak with an average signal of 6.7 and a peak width of ∼1.2 kb (Fig. 1E). An Mrr1 binding peak was similarly found upstream of the gene encoding Flu1 (Fig. 1F). The peaks associated with the *FLU1* ORF had an average signal of 4.8 and covered a length of ∼0.8 kb (Fig. 1F). The signal of the HF-Mrr1^Y813C^ peak upstream of the *MDR1* ORF was 1.5-and 2.1-fold higher than upstream of the *CDR1* and *FLU1* ORFs. Together, these data are consistent with previous reports of Mrr1 regulation of *MDR1* and provide evidence for direct regulation of *CDR1* and *FLU1* by Mrr1 in *C*. *lusitaniae*.

### Constitutive expression of *MDR1* reduces susceptibility to short chain azoles, while *CDR1* reduces susceptibility to long chain azoles

To investigate the phenotypic consequences of Mrr1 regulation of *MDR1*, *CDR1*, and *FLU1*, we determined the concentrations of various azoles required to inhibit 90% (MIC_90_) of the growth of strain U04 with Mrr1^Y813C^ and its *mdr1*Δ, *cdr1*Δ and *flu1*Δ derivatives. As the *C. albicans* homologs of Mdr1, Cdr1 and Flu1 were all capable of fluconazole (FLZ) efflux (29, 30, 39), we first evaluated the FLZ MIC_90_. The U04 strain expressing Mrr1^Y813C^ had a 32-fold higher FLZ MIC_90_ than the isogenic strain with Mrr1^ancestral^ (Fig. 2A & Table 1). The Mrr1^Y813C^ *mdr1*Δ mutant exhibited an 8-fold lower FLZ MIC_90_ than its parent *mrr1*Δ+*MRR1^Y813C^* strain (4 µg/ml vs 32 µg/ml; Fig. 2A & Table 1).

**Figure 2:**
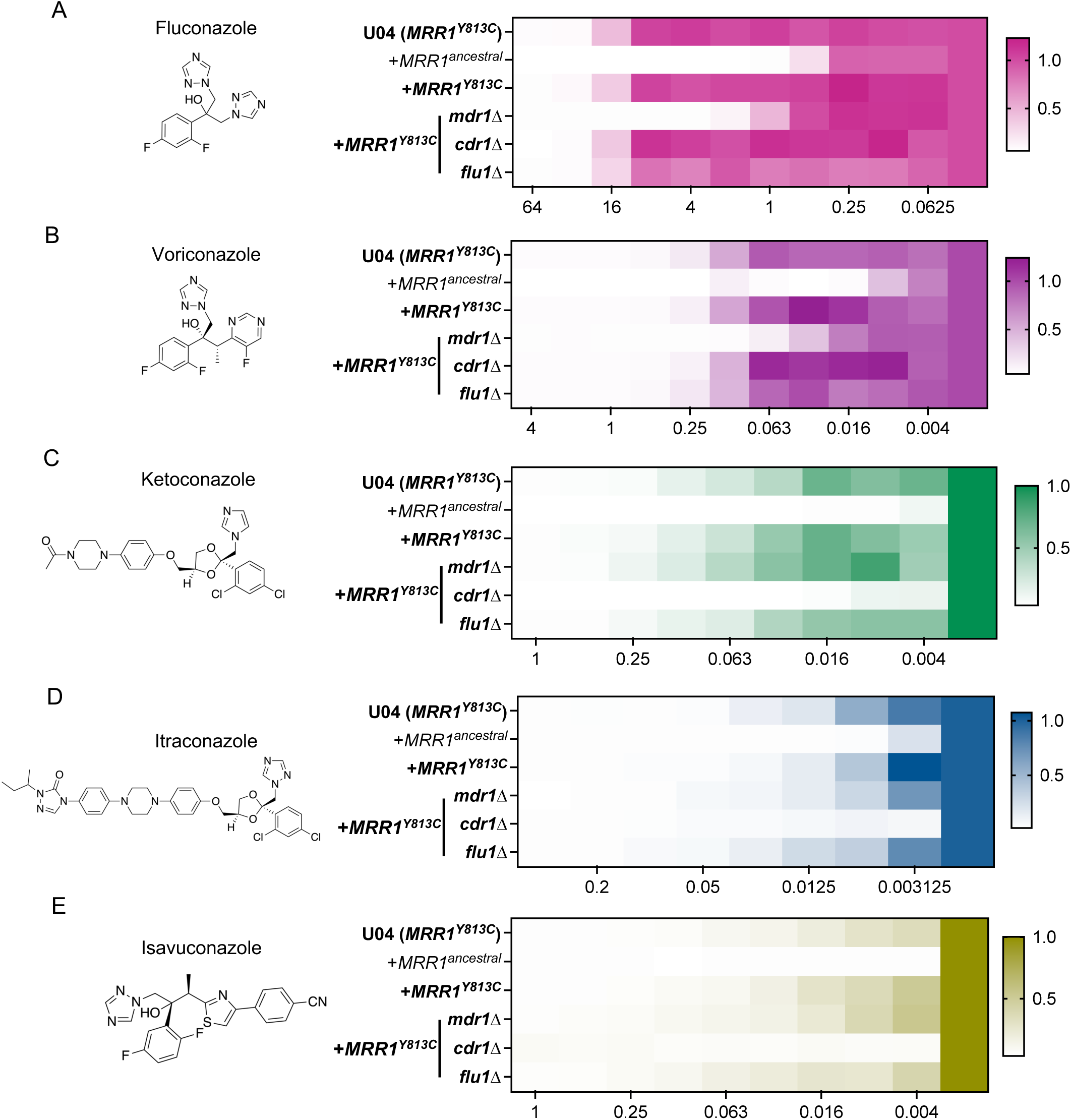
Effects of Mrr1 activity and *MDR1, CDR1* and *FLU1* on susceptibility to clinically relevant azoles. Structures and MICs (µg/ml) of Fluconazole (FLZ) (A), Voriconazole (VOR) (B), Ketoconazole (KTZ) (C), Itraconazole (ITZ) (D) and Isavuconazole (ISA) (E) are shown. MICs were determined using broth microdilution assays for strain U04 (native allele *MRR1^Y813C^*) and U04 mutants: *mrr1*Δ+*MRR1^ancestral^*, *mrr1*Δ+*MRR1^Y813C^* and *mdr1*, *cdr1* and *flu1* deletion mutants in the *mrr1*Δ+*MRR1^Y813C^* background. Heatmaps represent the optical density (600 nm) of the azole-treated wells normalized to the respective untreated strain controls (column on right). Drug concentrations in µg/ml are shown on the x-axis. The average from three independent experiments performed on different days is shown. Strains with constitutive Mrr1 activity are in bold.

**Table 1:** MIC_90_ of clinical azoles. MIC_90_ values were calculated from the broth microdilution assays in Figure 2. MIC_90_ was defined as the concentration at which 90% growth was inhibited. Fold differences in MIC_90_ relative to the azole-sensitive U04 *mrr1*Δ + *MRR1^ancestral^* are presented within parentheses (). FLZ – Fluconazole, VOR – Voriconazole, KTZ – Ketoconazole, ITZ – Itraconazole and ISA-Isavuconazole.

| Strains | FLZ | VOR | KTZ | ITR | ISA |
| --- | --- | --- | --- | --- | --- |
| U04 WT (*MRR1*<sup>Y813C</sup>) | 32 (32) | 0.5 (32) | 0.25 (32) | 0.05 (>16) | 0.0625 (>16) |
| U04 *mrr1*Δ + *MRR1*<sup>ancestral</sup> | 1 (1) | 0.0156 (1) | 0.0078 (1) | <0.003125 (1) | <0.0039 (1) |
| U04 *mrr1*Δ + *MRR1*<sup>Y813C</sup> | 32 (32) | 0.5 (32) | 0.25 (32) | 0.025 (>8) | 0.0625 (>16) |
| U04 *mrr1*Δ + *MRR1*<sup>Y813C</sup> *mdr1*Δ | 4 (4) | 0.125 (8) | 0.5 (64) | 0.025 (>8) | 0.125 (>32) |
| U04 *mrr1*Δ + *MRR1*<sup>Y813C</sup> *cdr1*Δ | 32 (32) | 0.5 (32) | 0.016 (2) | <0.003125 (1) | <0.0039 (1) |
| U04 *mrr1*Δ + *MRR1*<sup>Y813C</sup> *flu1*Δ | 32 (32) | 0.5 (32) | 0.25 (32) | 0.05 (>16) | 0.25 (>64) |

Although, the FLZ MIC_90_ values were unchanged in a *cdr1*Δ and *flu1*Δ mutant, the *flu1*Δ mutant grew slightly less well than the parent strain across concentrations (Fig. 2A & Table 1).

Similar Mdr1-dependent resistance was observed for the other short-tailed azole voriconazole (VOR) in strains with constitutively active Mrr1; the *mrr1*Δ+*MRR1^Y813C^* strain had a 32-fold higher VOR MIC_90_ than the *mrr1*Δ+*MRR1^ancestral^* strain (0.5 µg/ml vs 0.0156 µg/ml; Fig. 2B & Table 1). While the *mdr1*Δ mutation resulted in a 4-fold lower VOR MIC_90_ than the *mrr1*Δ+*MRR1^Y813C^* and the U04 WT (*MRR1^Y813C^*) strains, no difference in MIC_90_ was observed for the *cdr1*Δ and *flu1*Δ mutants (Fig. 2B & Table 1). Overall, strains expressing constitutively active Mrr1 exhibited similar Mdr1-mediated resistance to the triazoles FLZ and VOR. Interestingly, the *MDR1* deletion alone was not sufficient to abrogate resistance as the *mdr1*Δ mutant still had a 4-fold higher FLZ (4 µg/ml vs 1 µg/ml) and 8-fold higher VOR (0.125 µg/ml vs 0.0156 µg/ml) MIC_90_ values than the *mrr1*Δ+*MRR1^ancestral^*strain (Fig. 2A, B & Table 1) implying that there are redundant activities across other Mrr1-regulated azole resistance factors. Thus, our results suggest that Mdr1-mediates resistance to short-tailed azoles in strains with constitutive Mrr1 activity.

We evaluated the susceptibility of the different strains to the long-tailed azoles: ketoconazole (KTZ), itraconazole (ITR) and isavuconazole (ISA). The *mrr1*Δ+*MRR1^Y813C^* strain had a 32-, >8-and >16-fold higher MIC_90_ values for KTZ, ITR and ISA respectively, than the *mrr1*Δ+*MRR1^ancestral^* strain (Fig. 2C-E &Table 1). Consistent with prior reports of Cdr1-mediated resistance to long-tailed azoles (34, 40), the *cdr1Δ* strain had a >8-fold reduction in MIC_90_ values for KTZ, ITR and ISA than the *mrr1*Δ+*MRR1^Y813C^* parental strain (Fig. 2C-E & Table 1). The *mdr1*Δ and *flu1*Δ strains were not more susceptible to the tested long-tailed azoles than the parent strain (Fig. 2C-E & Table 1). These data indicate that constitutively active Mrr1 confers resistance to long-tailed azoles via Cdr1.

### Mrr1-regulated Mdr1 and Cdr1 decrease susceptibility to drugs from diverse classes

Transporter-mediated efflux of other antifungal compounds of agricultural and clinical relevance has been demonstrated (41) and strains with constitutive Mrr1 activity exhibited broad-spectrum resistance against multiple toxic substrates in a Biolog Phenotype Microarray screen (21). Thus, we evaluated the MICs of 5-flucytosine (5-FC), cycloheximide, myclobutanil, terbinafine and fluphenazine for *mrr1*Δ+*MRR1^ancestral^* and *mrr1*Δ+*MRR1^Y813C^* strains. Here on, MIC was defined as the concentration at which no visible growth was observed. The *mrr1*Δ+*MRR1^Y813C^*strain had a 2-to 32-fold increase in the MIC values of the different tested antifungals compared to the *mrr1*Δ+*MRR1^ancestral^* strain (Fig. 3A). Further, in the *mrr1*Δ+*MRR1^Y813C^* strain background, the *mdr1*Δ derivative resulted in increased susceptibility to 5-FC, cycloheximide and myclobutanil (Fig. 3B). The MIC values of cycloheximide and 5-FC decreased by 4-fold in the *mdr1*Δ mutant (Fig. 3B); support for Mdr1-mediated resistance against the pyrimidine analog 5-FC has been previously shown in *C. lusitaniae* (12, 25, 34). While the protein synthesis inhibitor cycloheximide was a substrate of both Mdr1 and Cdr1 (29) in *C. albicans*, the *cdr1Δ* mutation did not alter the cycloheximide resistance of the *mrr1*Δ+*MRR1^Y813C^*strain. For the agricultural triazole myclobutanil, the *mdr1*Δ and *cdr1*Δ mutants had 2-to 4-fold lower MIC values than the *mrr1*Δ+*MRR1^Y813C^* parental strain (2-4 µg/ml vs 8 µg/ml) (Fig. 3B). However, both still had 8-fold higher MIC values than the *mrr1*Δ+*MRR1^ancestral^* strain (2-4 µg/ml vs 0.25 µg/ml) suggesting that other Mrr1 targets contributed to myclobutanil resistance.

**Figure 3:**
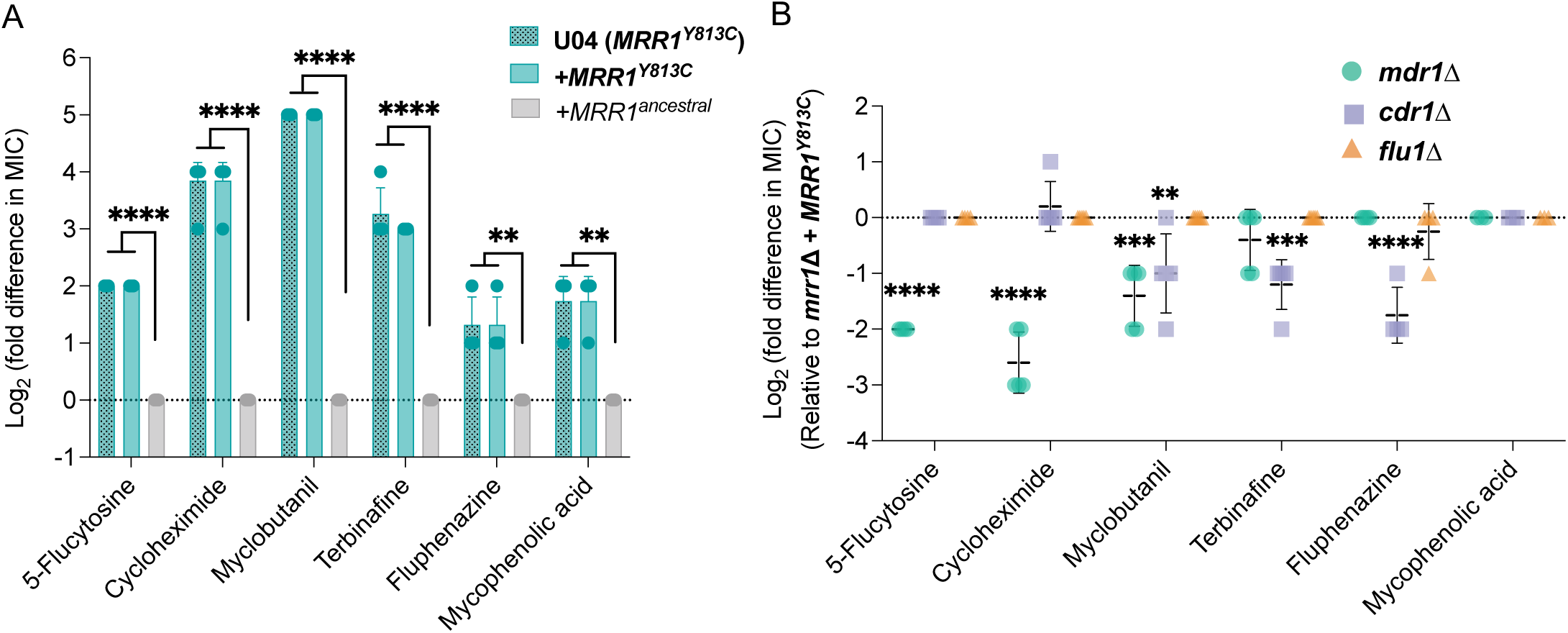
Effects of Mrr1 activity and *MDR1, CDR1* and *FLU1* on susceptibility to broad-spectrum antifungals. (A) Log_2_ transformed fold difference of MIC for diverse antifungals for strains U04 (native allele *MRR1^Y813C^*) and its *mrr1*Δ+*MRR1^ancestral^* and *mrr1*Δ+*MRR1^Y813C^*derivatives determined using broth microdilution assays. Data were normalized to that for *mrr1*Δ+*MRR1^ancestral^* strain. (B) Log_2_ transformed fold difference in MIC values of *mdr1Δ*, *cdr1Δ* and *flu1Δ* mutants normalized to their parent U04 *mrr1*Δ+*MRR1^Y813C^*. The data represent the mean ± SD from at least three independent experiments performed on different days. Strains with constitutive Mrr1 activity are in bold. Ordinary one-way ANOVA and Dunnett’s multiple comparisons testing with a single pooled variance were used to evaluate the statistical significance between strains for each antifungal. All significant comparisons are shown; *, p<0.05, **, p<0.01, ***, p<0.001 and ****, p<0.0001.

Susceptibility of other antifungals was dependent on Cdr1. The MICs for the allylamine antifungal terbinafine and the antipsychotic fluphenazine were lower in the *cdr1*Δ mutant. Since *FLU1* deletion made drug-sensitive *C. albicans* hypersusceptible to the metabolic inhibitor mycophenolic acid (MPA) (39), we also investigated the MPA susceptibility of our strains. Despite the *mrr1*Δ+*MRR1^Y813C^*having a 2-to 4-fold increase in MPA MIC relative to the *mrr1*Δ+*MRR1^ancestral^*strain, its MIC was not impacted by deletion of *FLU1*. The *mdr1*Δ and *cdr1*Δ mutants were also not more susceptible to MPA (Fig. 3B). Taken together, our results show that constitutive Mrr1 activity conferred resistance to a broad spectrum of antifungals, largely through its control of Mdr1-and Cdr1 with evidence for redundancy in Mrr1-regulated antifungal resistance mechanisms.

### Mrr1 directly regulates genes involved in diverse biological processes

To examine other genes that were co-regulated with *MDR1*, *CDR1* and *FLU1*, we identified additional genes that were differentially expressed due to a direct consequence of constitutive Mrr1 activity. There were twenty five genes that were differentially expressed when Mrr1 was constitutively active (Mrr1^Y813C^) compared to *mrr1*Δ and low-activity Mrr1 (21) (FDR < 0.05 and fold change ≥ 1.5) and had a HF-Mrr1^Y813C^ peak located within 1 kb from their ORF regions including *MDR1*, *CDR1*, and *FLU1* (Fig. 4A, Table S1A). These 25 genes will be referred to as the *C. lusitaniae* Mrr1 regulon (Table S1A). Slim Gene Ontology (GO) analysis of the *C. albicans* homologs of the *C. lusitaniae* Mrr1 regulon genes found transport, response to chemical, response to stress, and cellular homeostasis as the most enriched biological process terms (Table S1B). The Mrr1 regulon included two putative peptide transporters (*OPT1* and *OPT5*), two extracellular cell wall proteins (*ECM33* and *CSA1*), two involved in metal homeostasis (*CTR2* and *CFL4*), a putative glycerol transporter (*HGT10*/*STL1*), an alternative oxidase (*AOX2*), and multiple metabolic enzymes or putative oxidoreductases (Table S1A). Of note, the 77 indirect Mrr1 targets (Fig. 4A) were further enriched for transport, chemical and stress response processes in a Slim GO analysis of their *C. albicans* homologs (File S2A-B).

**Figure 4:**
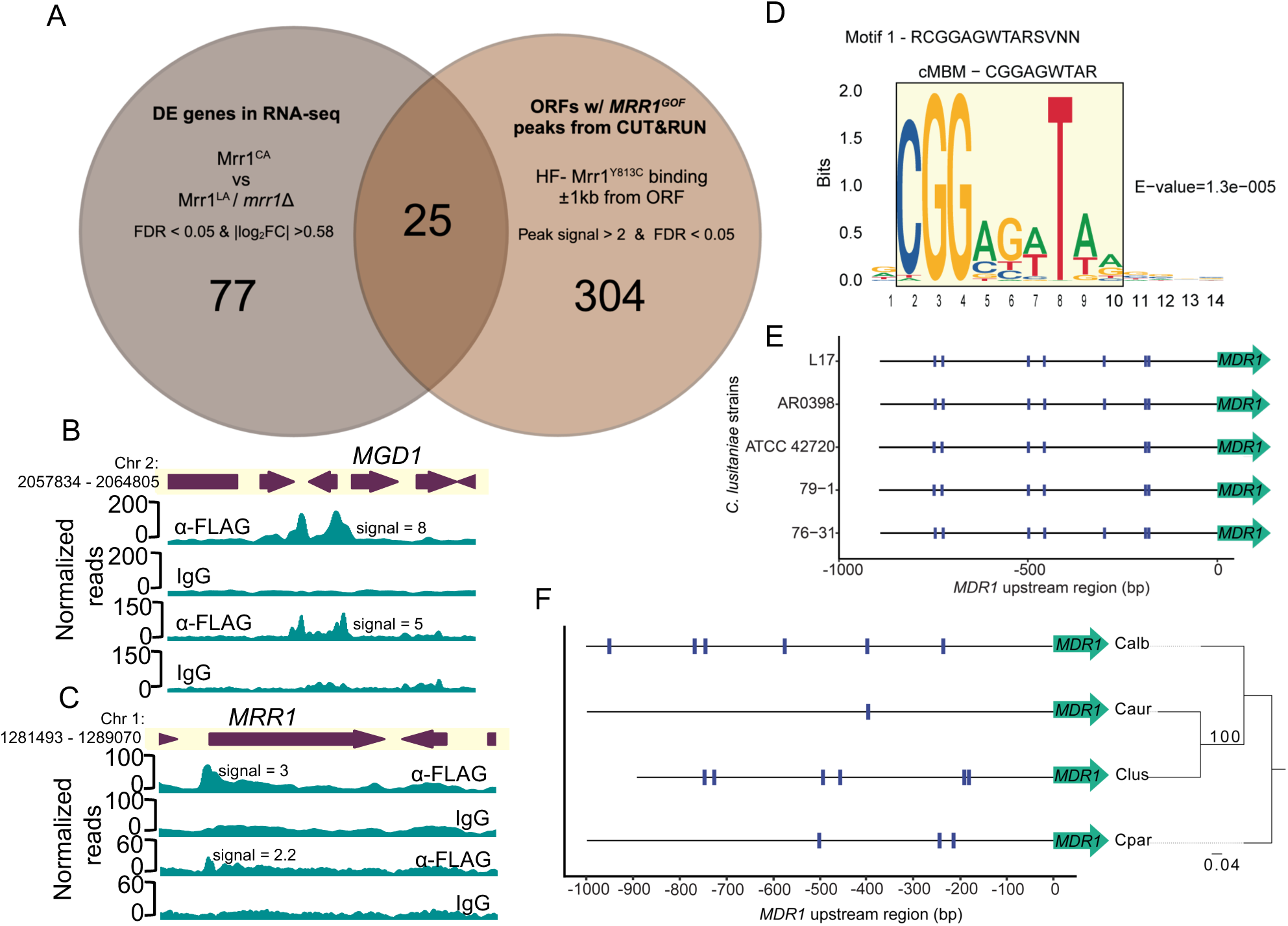
The Mrr1 regulon and the consensus Mrr1-binding DNA motif of *C. lusitaniae*. (A) Venn diagram shows the overlap between differentially expressed genes from RNA-seq in Demers *et al.* (21) and ORFs with HF-Mrr1^Y813C^ peaks in their intergenic regions from CUT&RUN. The 25 differentially regulated genes that have HF-Mrr1^Y813C^ peaks in either the 5’ or 3’ regions are listed in Table S1A. (B, C) HF-Mrr1^Y813C^ CUT&RUN read coverage plots normalized per 20 bp bin size. Chromosomal positions of regions containing *MGD1* and *MRR1* and adjacent genes are represented to scale with boxes and arrows. Peaks from HF-Mrr1^Y813C^-bound DNA recovered by an α-FLAG antibody and for the non-specific binding control recovered via IgG are shown. Signal indicates the average read density in α-FLAG relative to IgG within the peak region. Two independent experiments were performed and both are shown. (D) Sequence logo of the consensus motif detected within 100 bp of CUT&RUN peak summits by STREME. E-value is an estimate of motif significance. The 9-nt <u>c</u>onsensus <u>M</u>rr1-<u>b</u>inding motif (cMBM) is boxed in yellow. (E) cMBM location (blue hatches) in the ∼890 bp upstream intergenic regions of *MDR1* in different *C. lusitaniae* strains. (F) cMBM location (blue hatches) in the 1 kb upstream intergenic regions of the *MDR1* homologs of *C. parapsilosis* CDC317, *C. auris* B11205*, C. albicans* SC5314 and *C. lusitaniae* ATCC 42720. The *C. lusitaniae* ATCC 42720 upstream intergenic region is 893 bp. The phylogenetic tree was constructed using the *MDR1* nucleotide sequences.

We previously showed that the *C. lusitaniae* Mrr1 induced *MGD1* and *MGD2* in the presence of exogenous MGO (24), a toxic 2-oxo-aldehyde released by metabolically dysregulated cells and activated macrophages at sites of infection (42). Further, upregulation of *MGD1* and *MGD2* by constitutively active Mrr1 conferred a growth advantage in the presence of MGO (24). Methylglyoxal dehydrogenases are co-regulated with *MDR1* in several other *Candida* spp. including *C. albicans* and *C. auris* (18, 21, 23, 25, 26, 28). Interestingly, despite high expression of both *MGD1* and *MGD2* transcripts in strains with Mrr1^Y813C^ (21), only the promoter regions of *MGD1* had a HF-Mrr1^Y813C^ CUT&RUN peak with an average signal of 6.5 (Fig. 4B). A HF-Mrr1^Y813C^ peak of average signal 2.8 was also present in the promoter regions of *MRR1* (Fig. 4C) indicating at a mechanism for potential positive self-regulation of *MRR1* transcripts which is consistent with previously published RNA-seq data (21). Three Mrr1 regulon genes (*CLUG_04865, CLUG_01574,* and *CLUG_04429*) had no clear homologs in *C. albicans*, but did have homologs in the more closely related *C. auris*. Although not differentially expressed in the U04 transcriptome, a putative alcohol dehydrogenase *(CLUG_00171*) and a putative phospholipase C (*CLUG_01152*) had HF-Mrr1^Y813C^ peaks in their promoter regions and were upregulated in the clinical *C. lusitaniae* P3 isolate with a *MRR1^V668G^* GOF allele (25). Five genes were less abundant in strains with activated Mrr1 (Table S1A); one of these, *CLUG*_*01020* (*STL1*), was the only locus with an HF-Mrr1^Y813C^ peak in its 1 kb downstream intergenic region with no peak in its upstream region (Table S1A).

### Definition of an Mrr1-binding DNA motif that is conserved across species

To better understand direct Mrr1 regulation of targets, we used the STREME algorithm (43) to determine if specific motifs were enriched within sequences corresponding to 329 HF-Mrr1^Y813C^ CUT&RUN peaks (File S1). The 100 bp sequences upstream and downstream of peak summits (the most enriched point within an identified peak) were used as input for discriminative *de novo* motif discovery (43). A set of sequences chosen at random from *C. lusitaniae* L17 genome and matched in length and number was used as background to identify enriched motifs in the input set. A 14-nucleotide (nt) consensus sequence RCGGAGWTARSVNN was the topmost motif predicted by STREME (Fig. 4D).

When this consensus sequence was scanned for in the upstream intergenic regions of *MDR1* from *C. lusitaniae* L17 and ATCC 42720 (ASM383v1), the motif was observed seven times in the *C. lusitaniae* L17 *MDR1* promoter region and six of these were conserved in *C. lusitaniae* strain ATCC 42720 (Fig. S1A). The 14-nt consensus sequence had substantial nucleotide ambiguity at both ends (positions 1 and 11-14) (Fig. 4D). Therefore, for subsequent motif analyses, we focused on the internal 9-nt CGGAGWTAR motif (Fig. 4D, boxed). The 9-nt CGGAGWTAR motif and the 14-nt RCGGAGWTARSVNN motif were similarly detected in the promoter regions of *MDR1* in both strains (Fig. S1A). Henceforth, we refer to the 9-nt CGGAGWTAR motif as the <u>c</u>onsensus <u>M</u>rr1-<u>b</u>inding DNA <u>m</u>otif (cMBM) (Fig. 4D)

The analysis of *MDR1* promoter sequences from the clinical isolate AR0398 (GCA_032599225.1) and two distantly related environmental isolates 79-1 (GCA_032599145.1) and 76-31 (GCA_032599085.1) found the six cMBM sites detected in strains L17 and ATCC 42720 (44). Each of the cMBMs were at identical positions and orientations relative to the *MDR1* translational start sites across the different strains (Fig. 4E). cMBMs were also found upstream of *CDR1* and they were again conserved in position in both L17 and ATCC 42720 strains despite differences in the length of the *CDR1* adjacent intergenic regions (Fig. S1B). At least one cMBM, and often multiple cMBMs, were found within the peak spanning regions associated with all but two of the genes in the Mrr1-regulon (File S3).

We also scanned for the cMBM in the promoter sequences of the *MDR1* and *CDR1* homologs in *Candida* spp. At least three copies of cMBM were found in the *MDR1* and *CDR1* promoter sequences of *C. albicans* and *C. parapsilosis*, and one cMBM in *C. auris* (Fig. 4F & S1C). In the case of *C. albicans*, two cMBMs occurred in locations previously annotated to be important for *MDR1* transcriptional regulation. These cMBMs were discovered between the-200 to-400 regions which encompassed the benomyl response element (-260 and-296) (45) and the Mrr1-binding region that contained the *C. albicans* Mrr1-binding DNA motif DCSGHD (-342 to-492) (38). In a ChIP-qRT analysis of Mrr1 binding to the *C. albicans MDR1* promoter, DNA recovery was highest at these cMBM-containing regions relative to the rest of the *MDR1* promoter sequence (20). Together these data strongly suggest that the consensus Mrr1-binding DNA motif discovered in *C. lusitaniae* is conserved in other *Candida* species. Transcription factors of the zinc-cluster family, which includes Mrr1, typically bind to CGG motifs occurring as direct, inverted or everted repeats (46).

### Constitutively active and low-activity Mrr1 localize to similar genomic regions in *C. lusitaniae*

Previous studies on *C. lusitaniae* Mrr1 suggested that expression at some loci (e.g. *MDR1* and *MGD1*) (21, 24, 25) was repressed by low-activity Mrr1 variants and induced in the presence of benomyl and MGO inducers of Mrr1 or by constitutively activate Mrr1 variants. Thus, we compared DNA localization of the HF-Mrr1^Y813C^ to the genome-wide binding of low activity HF-Mrr1^ancestral^ in the absence of Mrr1 inducing stimuli. Using the same parameters as for the analysis of HF-Mrr1^Y813C^, we found around 1,276 peaks associated with HF-Mrr1^ancestral^–bound DNA (File S4). The *MDR1* intergenic region revealed a significant HF-Mrr1^ancestral^ peak that spanned a region of ∼1.6 kb and had a signal of 15.1 (Fig. 5A). HF-Mrr1^ancestral^ peaks were also found upstream of *CDR1* and *FLU1* (1.6 and 2.2 peak signal, respectively; Fig. 5B-C). Comparison of HF-Mrr1^ancestral^ and HF-Mrr1^Y813C^-bound sites upstream of *MDR1*, *CDR1* and *FLU1* exhibited a striking similarity in their peak profiles (Fig. S2A-C). The remarkable overlap of HF-Mrr1^ancestral^ and HF-Mrr1^Y813C^ CUT&RUN peaks present in over 930 genomic locations (Fig. 5D) suggest that Mrr1-mediated repression and induction are not due to differences in Mrr1 localization to the DNA.

**Figure 5:**
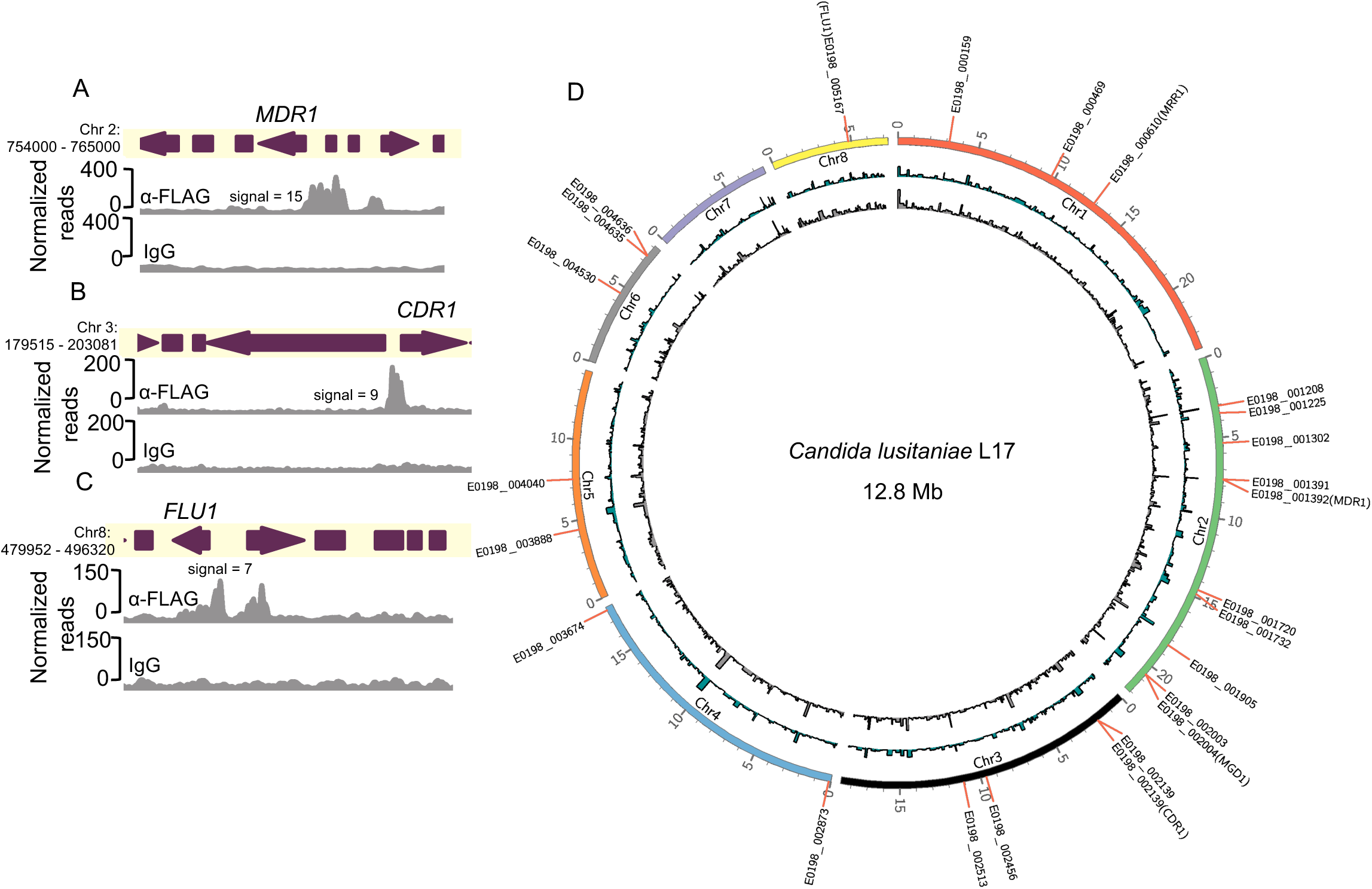
Local and global binding profiles of constitutively active and low-activity Mrr1. (A-C) HF-Mrr1^ancestral^ CUT&RUN read coverage plots normalized per 20 bp bin size. Chromosomal positions of regions containing *MDR1, CDR1* and *FLU1* and adjacent genes are represented to scale with boxes and arrows. Peaks from HF-Mrr1^ancestral^-bound DNA recovered by an α-FLAG antibody and for the non-specific binding control recovered via IgG are shown. Signal indicates the average read density in α-FLAG relative to IgG within the peak region. (D) Circos plot showing global CUT&RUN-determined Mrr1-binding peaks of HF-Mrr1^Y813C^ (in blue) and HF-Mrr1^ancestral^ (in grey) in the *C. lusitaniae* L17 genome. Mrr1-binding peaks with a signal ≥2-fold compared to their respective IgG backgrounds and up to 1 kb away from the nearest ORF from Experiment 1 were used (see Supplemental Files 1 and 4). The genomic positions of the 25 differentially expressed genes that constitute the Mrr1-regulon are marked with the L17 gene IDs.

In Demers *et al.*(21), we characterized *MRR1* alleles with GOF mutations that resulted in constitutive activity and Mdr1-dependent FLZ-resistance (Fig. 6A) as well as alleles with both GOF mutations and secondary suppressor mutations that restored the inducible low activity state such as *MRR1^L1191H+Q1197*(L1Q1*)^* (Fig. 6A). The *mrr1*Δ+*MRR1 ^L1Q1*^* strain had more than a 32-fold lower FLZ MIC value (0.125 µg/ml vs 32 µg/ml) than strains with *MRR1^GOF^* alleles (*MRR1^Y813C^* and *MRR1^L1191H^*) (Fig. 6B). Since GOF mutations in Mrr1 did not affect DNA localization, we evaluated whether secondary suppressor mutation(s) altered these interactions by performing CUT&RUN on U04 *mrr1*Δ strains expressing HF-Mrr1^L1Q1*^ from its endogenous promoter. Western blot confirmed that the truncated HF-Mrr1^L1Q1*^ was present at levels similar to that of the full-length HF-Mrr1^ancestral^ and HF-Mrr1^Y813C^ (Fig. S3A). The HF-tag did not modify Mrr1^L1Q1*^ activity as strains expressing tagged Mrr1^L1Q1*^ exhibited similar 32- to 64-fold lower FLZ MIC as untagged Mrr1^L1Q1*^ when compared to strains expressing the constitutively active Mrr1^Y813C^ variant (Fig. S3B). Our CUT&RUN analysis found HF-Mrr1^L1Q1*^- bound DNA to be significantly enriched in the upstream intergenic regions of *MDR1*, *CDR1* and *FLU1* ORFs (Fig. 6C-E) with a peak profile identical to HF-Mrr1^ancestral^ and HF-Mrr1^Y813C^. The HF-Mrr1^L1Q1*^ peak recapitulated the 1.5- and 2-fold higher signal upstream of *MDR1* relative to *CDR1* and *FLU1*. Across the entire *C. lusitaniae* genome, the HF-Mrr1^L1Q1*^- bound genomic sites (see File S5 for peaks) were strikingly similar to the HF-Mrr1^ancestral^ and HF-Mrr1^Y813C^-bound sites suggesting that secondary suppressor mutation(s) do not likely impact Mrr1 localization to the DNA (Fig. S4). Hence, our results illustrate that Mrr1 localization at the *C. lusitaniae* DNA are unaltered by the tested mutations and are independent of Mrr1 activation state.

**Figure 6:**
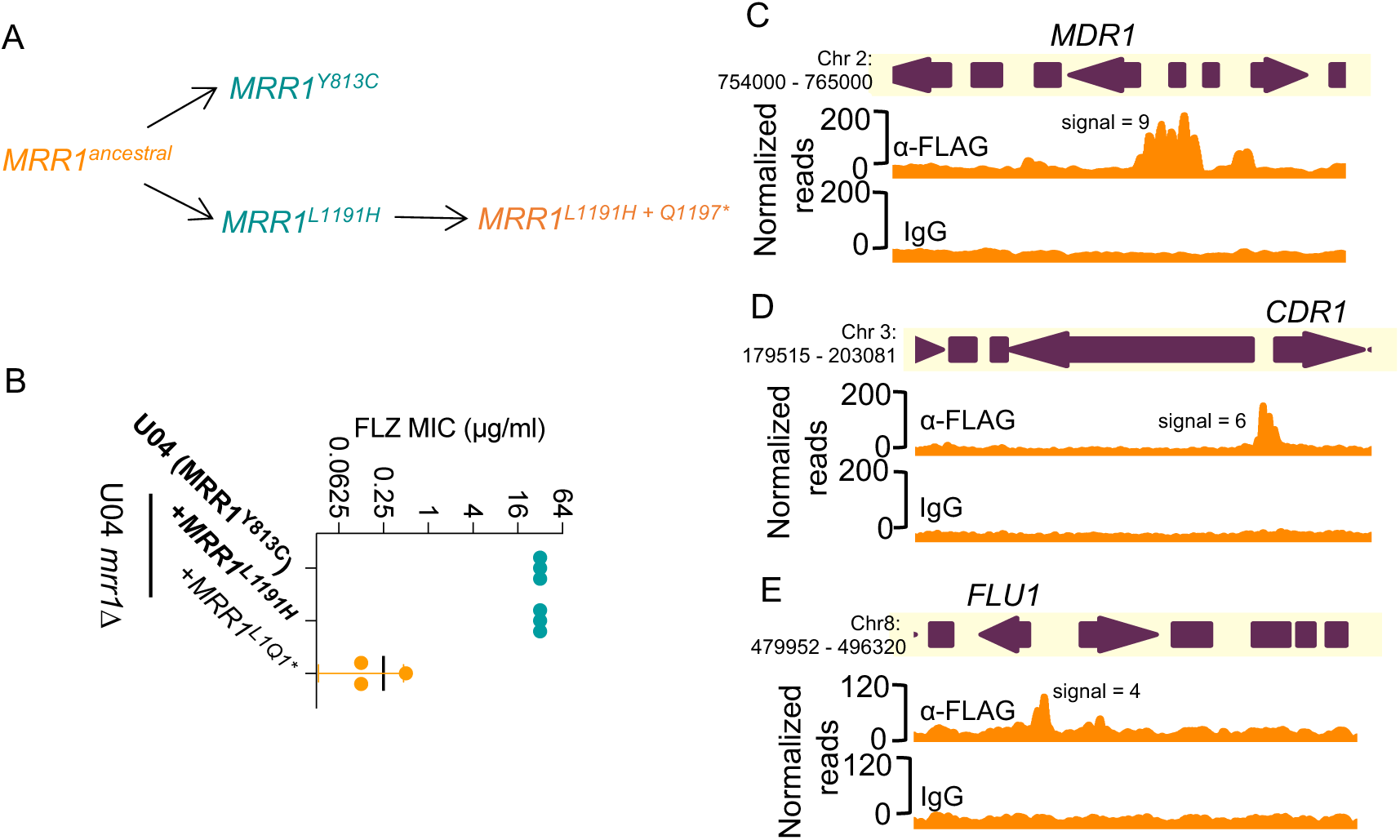
Evolution of naturally acquired *MRR1* mutations and binding profiles of low-activity Mrr1. (A) Schematic of the clinically evolved *MRR1* alleles reported in Demers *et al.* (21). The asterisk indicates a nonsense mutation. Alleles in blue and orange encode for constitutively active and low-activity Mrr1 variants, respectively. (B) FLZ MIC of U04 clinical isolate (native allele *MRR1^Y813C^*) and U04 *mrr1*Δ complemented with *MRR1^L1191H^* or *MRR1^L1Q1*^ ^(L1191H^ ^+ Q1197*)^* was determined by broth microdilution assays. The data shown represent the mean ± SD from three independent experiments. Strains with constitutive Mrr1 activity are in bold. (C-E) HF-Mrr1^L1Q1*^ CUT&RUN read coverage plots normalized per 20 bp bin size. Chromosomal positions of regions containing *MDR1, CDR1* and *FLU1* and adjacent genes are represented to scale with boxes and arrows. Peaks from HF-Mrr1^L1Q1*^-bound DNA recovered by an α-FLAG antibody and for the non-specific binding control recovered via IgG are shown. Signal indicates the average read density in α-FLAG relative to IgG within the peak region.

## DISCUSSION

In this study, we demonstrated that constitutively active *C. lusitaniae* Mrr1 directly upregulates several multi-drug transporter-encoding genes, including *MDR1* and *CDR1,* leading to reduced susceptibility to both short-tailed and long-tailed azoles and other antifungals (Fig. 1-3). The coordinated regulation of both *MDR1* and *CDR1* by Mrr1 in *C. luistaniae* differs from their regulation in the well-studied species *C. albicans* wherein Mrr1 is the primary regulator of *MDR1* and Tac1 is the main CDR1 transcriptional activator (23, 33). We identified a consensus Mrr1 binding motif (cMBM; CGGAGWTAR) that colocalized with Mrr1 CUT&RUN peaks and that was present in multiple positions within the peaks in regions adjacent to *C. lusitaniae MDR1*, *CDR1* and in almost all other Mrr1-regulated genes (Fig. 4D & File S2). The cMBM sequences in Mrr1 peak regions were conserved in other *C. lusitaniae* strains (Fig. 4E & S1B). Furthermore, the cMBM was also enriched in the regions upstream of *MDR1* homologs in *C. albicans*, *C. auris* and *C. parapsilosis,* and in *C. albicans*, the cMBM was present in regions shown to bind *C. albicans* Mrr1 (20). The cMBM was also upstream of *C. lusitaniae* and *C. parapsilosis CDR1* which is consistent with reports that constitutive Mrr1 activity also induces expression of *CDR1* in these species. Furthermore, we noted the presence of cMBMs in regions upstream of *CDR1* in species that have no reports for Mrr1 regulation of *CDR1* including *C. albicans* and *C. auris* (33, 47, 48). Consistent with the potential for Mrr1 regulation of *CDR1* in *C. albicans*, a ChIP-ChIP analysis detected Mrr1 in the upstream regions of *CDR1* (49), though *CDR1* was not reported as an Mrr1 target because its expression was not increased by constitutively active Mrr1.

Studies in *C. albicans* and recent work in *C. auris* have shown Tac1 with an activating mutation upregulates *CDR1* expression and the *C. albicans* Tac1 regulates *CDR1* by binding a consensus CGGN_4_CGG motif in the promoter region (49). Though *C. lusitaniae* has a Tac1 homolog (Clug_02369) (34) and a CGGN_4_CGG motif at-761 in the *CDR1* promoter region (data not shown), strains with low Mrr1 activity and a *cdr1*Δ mutant had similar susceptibilities (MIC 12.5 – 25 µg/ml) to the Cdr1 substrate fluphenazine (Fig. 3) (22, 50). In fact, while activating mutations in *TAC1* have been characterized in FLZ-resistant *C. parapsilosis* (51) and *C. auris* (52), to our knowledge, there are no reports on activating mutations in the *TAC1* gene leading to FLZ resistance in *C. lusitaniae*. While we (21) and others (34) have shown that Mrr1 is sufficient to upregulate *C. lusitaniae CDR1*, Tac1 may induce *CDR1* under conditions not tested in this study. For instance, estradiol is an inducer of Tac1-mediated *CDR1* expression in *C. albicans* (53, 54).Together, these data underscore the evolutionary plasticity in transporter regulation in *Candida* spp. through the adoption of targets from one transcriptional circuit to another (Fig. 7A; (55–57)). In the case of *C. lusitaniae*, the coordinated regulation of drug efflux proteins may be a mechanism for cross-resistance to multiple antifungals and may promote the development of other resistance mutations through a reduction in drug susceptibility.

**Figure 7:**
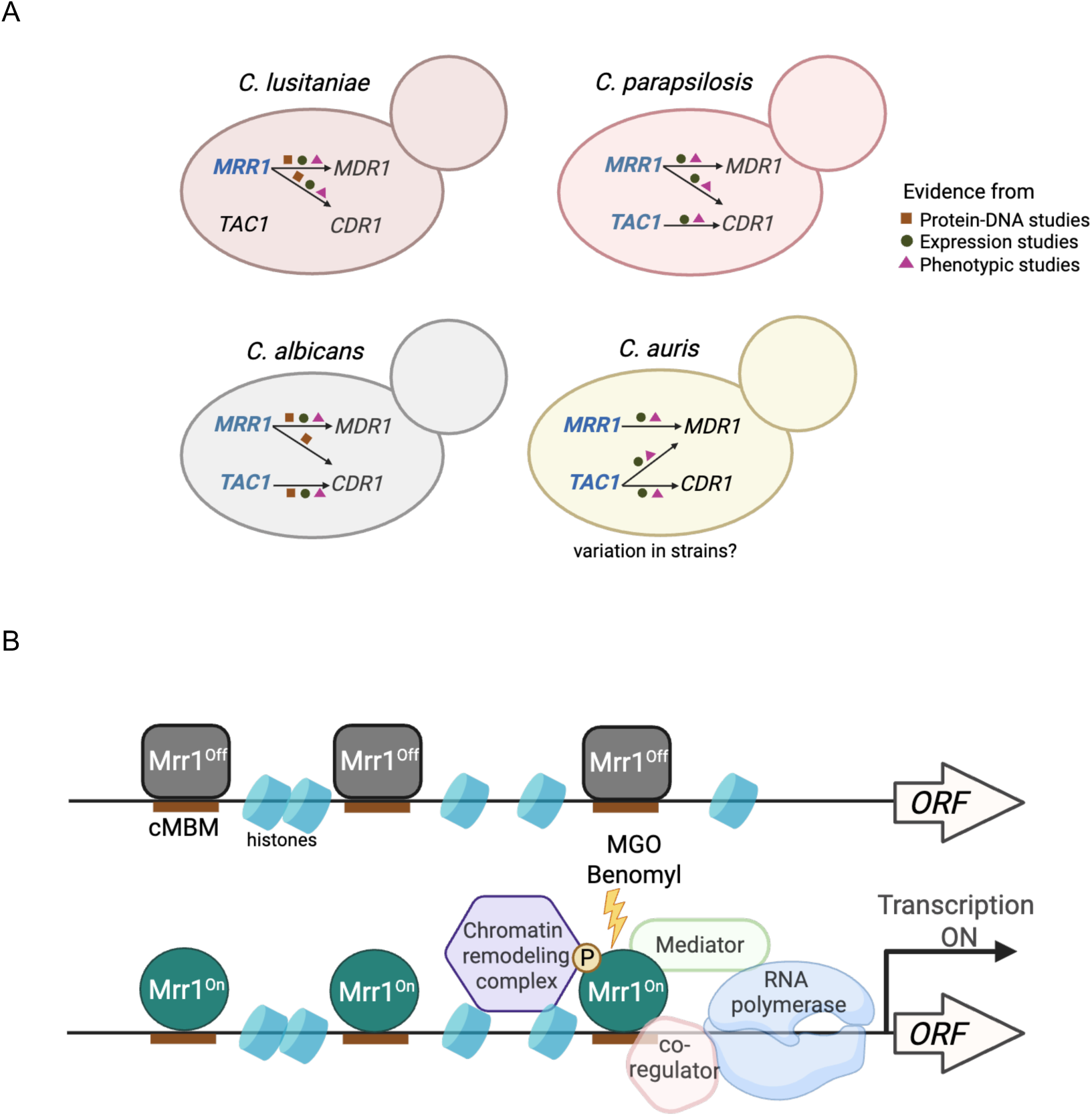
Models for Mrr1 regulation in *Candida* spp.. (A) Evolution of transporter regulation in *Candida* spp. Genes having characterized GOF mutations are in blue. The shapes indicate the type of experimental data used to support the model including protein-DNA studies (this study; 20, 38, 49), expression and phenotypic studies (18–21, 23–28, 32–35, 40, 47, 48, 50–52). (B) Possible mechanism(s) for gene induction by Mrr1 based on published studies (20, 38, 46, 54, 58–60). The mechanisms that impact Mrr1-mediated gene expression may vary between promoters and conditions within a strain, and there may be differences across strains and species.

Our data on Mrr1 levels and Mrr1 variants binding to upstream, or in some cases downstream, regions of Mrr1-regulated genes provide insight into Mrr1 regulation. First, we found that Mrr1 variants with differing activities did not have differences in total protein levels (Fig. S3A). Second, activated Mrr1 and inducible but inactive Mrr1 had indistinguishable localization at all cMBMs (Fig. S4) which is consistent with ChIP-qRT analysis of Mrr1 interactions with the *MDR1* promoter region in *C. albicans* (20). Third, the subset of genes repressed by inducible but inactive Mrr1 had similar Mrr1 localization in their promoter regions as those genes that were not repressed by Mrr1 (Fig. S2A). Thus, Mrr1 is likely regulated through mechanisms such as induced conformational change by ligand, co-factor binding (58), phosphorylation (59) or differential activity of co-regulatory proteins. These mechanisms are not mutually exclusive (46). In *C. albicans*, changes in activity of coregulatory proteins (Cap1 or Mcm1 (38, 60)), mediator (20, 54) or chromatin remodeling complexes like the Swi/Snf complex influence Mrr1 induction of *MDR1* and other genes (20) (Fig. 7B). The >1 kb width of our CUT&RUN peaks is consistent with the presence of multiple cMBMs in regions adjacent to Mrr1-regulated genes and may also reflect the presence of co-regulators or chromatin remodeling complexes that could influence micrococcal nuclease access to DNA. The involvement of multiple regulatory mechanisms allows for the controlled and differential expression of unique gene subsets in different strain backgrounds (12) in response to environmental cues that may be present in an infection environment (e.g. decreased nutrient availability or metabolites like methylglyoxal or inflammatory molecules) The presence of diverse regulatory mechanisms may promote survival under diverse conditions and may also promote the evolution of novel regulatory circuits across species and even strains (Fig. 7A).

For many azoles, there was an 8- to 16-fold increase in the MIC values of strains with activated Mrr1 variants compared to strains with low-activity Mrr1 and these differences were dependent on either Mdr1 (FLZ and VOR) or Cdr1 (KTZ, ITR and ISA) (Fig. 2 & Table 1). In the *C. lusitaniae* clinical isolate P3 which has activated Mrr1, the *mdr1*Δ*cdr1*Δ double mutant was even more susceptible to azoles (FLZ, VOR, ITR) than their single deletion mutants (34). Redundancy in transporter efflux was also observed in the case of other broad-spectrum antifungals. In addition to Mdr1 and Cdr1, susceptibility to the tested antifungals could also be mediated by other efflux pumps in the Mrr1 regulon including the MFS family transporter Flu1. While *FLU1* is a conserved Mrr1 target in other *Candida* spp. such as *C. albicans* (17) and *C. parapsilosis* (35), the promiscuity for substrates between transporters may have concealed any apparent contribution of Flu1 to efflux in a *MDR1*/*CDR1* overexpression strain (Fig. 2, 3 & Table 1). Beyond drug efflux, the *C. lusitaniae* Mrr1 regulon (Table S1A) included genes involved in other transporter activities like oligopeptide transport (*OPT1*), and chemical and stress response which is consistent with published Mrr1 regulons of *C. albicans* (23) and *C. parapsilosis* (28, 35). In *C. auris*, the *OPT1* homolog is upregulated in response to stress such as antifungal exposure (61) or macrophage phagocytosis (62). While *OPT1* may be involved in nutrient uptake under stress conditions (63), other metabolic factors including aldehyde and methylglyoxal dehydrogenases and aldo-keto reductases are speculated to protect cells from reactive molecules generated by azole stress (29). Thus, the Mrr1-regulated metabolic and stress response genes may be important for the persistence of the *MRR1^GOF^* mutants *in vivo* or could lead to the selection for *MRR1^GOF^* mutants in drugless conditions (18). Understanding Mrr1 regulation of these additional targets across *Candida* spp. can provide insights into the mechanisms that change multi-drug transporter regulation in *Candida*.

## Materials and Methods

### Strains and growth conditions

Strains used in this study are listed in Table S2. All strains were stored as frozen stocks with 25% glycerol at-80°C and maintained regularly on YPD (1% yeast extract, 2% peptone, 2% glucose, 1.5% agar) plates incubated at 30°C then stored at room temperature. Strains were grown in YPD liquid medium (5 ml) at 30°C on a roller drum for ∼16 h prior to inoculation into specified culture conditions. For drug susceptibility assays, cells were grown in RPMI-1640 (Sigma, containing L-glutamine, 165 mM MOPS, 2% glucose, pH 7) liquid, as noted. *Escherichia coli* strains were grown in LB with either 150 µg/ml carbenicillin or 15 µg/ml gentamicin as necessary to maintain plasmids.

### Strain construction

Gene replacement constructs for knocking out *MRR1* (*CLUG_00542*, as annotated in (18)) and *MDR1* (*CLUG_01938/9* (18)) were generated by fusion PCR, as described in Grahl *et al.* (64). All primers (IDT) used are listed in Table S3. Briefly, 0.5 to 1.0 kb of the 5’ and 3’ regions flanking the gene was amplified from U04 DNA, isolated using the MasterPure Yeast DNA Purification Kit (epiCentre). The nourseothricin (*NAT1*) or hygromycin B (HygB) resistance cassette was amplified from plasmids pNAT (65) and pYM70 (66) respectively. Nested primers within the amplified flanking regions were used to stitch the flanks and resistance cassette together. Gene replacement constructs for knocking out *CDR1* (*CLUG_03113*) and *FLU1* (*CLUG_05825*) were generated by introducing 30- to 50-bp of the 5’ and 3’ regions flanking the gene of interest into the replacement *NAT1* cassette using PCR. PCR products for transformation were purified and concentrated with the Zymo DNA Clean & Concentrator kit (Zymo Research) with a final elution in molecular biology grade water (Corning).

### Plasmids for complementation of *MRR1*

Plasmids for complementing untagged *MRR1* were created as described in Biermann *et al.* (24). Plasmids for complementing N-terminal HF-tagged *MRR1* were made as follows. We amplified i) the 6xHis-3xFLAG-tag from an *HF-MRR1* tagged *C. albicans* strain DH2561 using primers ED207 and ED208, ii) the ∼1150 bp upstream region of the *MRR1* gene for homology, from the respective *MRR1* allele complementation plasmids, using primers ED103 and ED206 and iii) ∼1500 bp of the *MRR1* gene using primers ED209 and ED132. The 6xHis-3xFLAG-tag is placed after the first codon of *MRR1*. PCR products were cleaned up using the Zymo DNA Clean & Concentrator kit (Zymo Research). The amplified PCR products were assembled into a pMQ30 vector using the *S. cerevisiae* recombination technique described in Shanks *et al.* (67). Plasmids created in *S. cerevisiae* were isolated using a yeast plasmid miniprep kit (Zymo Research) and transformed into High-Efficiency NEB®5-alpha competent *E. coli* (New England BioLabs). *E. coli* containing pMQ30-derived plasmids were selected for on LB containing 15 µg/ml gentamicin. Plasmids from *E. coli* were isolated using a Zyppy Plasmid Miniprep kit (Zymo Research) and subsequently verified by Sanger sequencing. *MRR1* complementation plasmids were linearized with the NotI-HF restriction enzyme (New England BioLabs), cleaned up using the Zymo DNA Clean & Concentrator kit (Zymo Research) and eluted in molecular biology grade water (Corning) before transformation of 2 µg into *C. lusitaniae* strain U04 *mrr1*Δ as described below.

### Strain construction

Mutants were constructed as previously described in Grahl *et al.* using an expression-free ribonucleoprotein CRISPR-Cas9 method (64). One to 2 µg of DNA for gene knockout constructs generated by PCR or 2 µg of digested plasmid, purified and concentrated with a final elution in molecular biology grade water (Corning), was used per transformation. *E. coli* strains containing the complementation and knockout constructs and crRNAs are listed in Table S2 and Table S3 respectively. Transformants were selected on YPD agar containing 200 µg/ml nourseothricin or 600 µg/ml hygromycin B.

Mutants for *CDR1* and *FLU1* were generated using a microhomology-mediated end-joining (MMEJ) repair method as described in Al Abdallah *et al.* (68). One to 2 µg of DNA for gene knockout constructs generated by PCR were used for transformation. crRNAs (IDT) used to target the 5’ and 3’ end of the gene of interest are listed in Table S3. *CDR1* and *FLU1* knockout transformants were selected on YPD agar containing 200 µg/ml nourseothricin.

### Protein Isolation

Overnight cultures were back diluted into 50ml YPD and grown to exponential phase (∼5 h) at 30°C. Harvested cells were snap-frozen using ethanol and dry ice and stored at-80°C. Thawed cell pellets were resuspended in a homogenization buffer (10 mM Tris-HCl, 150 mM NaCl, and 5 mM EDTA, adjusted to pH 7.4 and 10% sucrose) with protease inhibitor (2x Halt protease, Thermo Scientific) and mixed with an equal volume of 1:1 of 0.5- and 1-mm silica bead mix in a bead beating tube (VWR). Bead beating was carried out for six cycles at a speed of 5.65 for 20 seconds, with a one-minute rest on ice between each cycle. The top liquid phase was collected and centrifuged to remove any cell debris. Supernatants were transferred to new tubes and stored at-80°C. Protein concentrations were determined using the Bradford assay (Quick Start Bio-Rad) with a standard curve generated using serial dilutions of 2 mg/ml bovine serum albumin (BSA).

### Western Blot for HF-Mrr1 detection

Samples were diluted to equal concentrations in sample buffer (3.78% Tris, 5% SDS, 25% sucrose at pH 6.8 and 0.04% bromophenol blue prepared as a 5x stock solution. β-mercaptoethanol (0.05%) was freshly added). Samples were heated for 10 min at 95°C and loaded into 6.5% SDS page gels along with a BioRad All Blue Precision Plus MW marker. The gel was run for ∼40 min at 180 V. The BioRad Turboblot semi-dry transfer system with custom settings (1.3A constant and 25V for 15 min) was used to transfer the protein bands to a LF-PVDF membrane (Immobilon Product IPFL00010). The blots were processed using the standard LICOR protocol for Western blotting, including the optional drying step after transfer and REVERT total protein staining. A milk-based blocking buffer was used instead of the Odyssey blocking buffer. The α-FLAG monoclonal antibody (1mg/mL) (Sigma-Aldrich M2 or ThermoFisher FG4R) was diluted 3000-fold in blocking buffer with 0.1% Tween-20. The goat α-mouse secondary antibody (1 mg/mL) labelled with IRDye 700CW was diluted 15,000-fold in blocking buffer with 0.1% Tween-20. Blots were imaged using an Odyssey CLX scanner (LICOR) and analyzed using the Empiria software (LICOR).

### CUT&RUN experimental setup and sequencing

Overnight cultures were back diluted into 50 ml YPD and grown to exponential phase (∼5 h) at 30°C. Samples were processed using the Epicypher CUT&RUN kit (Epicypher) as per the protocol described in Qasim *et al*. (69). Briefly, yeast nuclei were isolated from the thawed cell pellets using Zymolase 100T (Zymoresearch). Digitonin (0.01%) was added to all buffers used hereafter to permeabilize nuclei and prevent bead clumping. The isolated nuclei were bound to activated concanavalin A (ConA)-coated magnetic beads. The nuclei-bound ConA-beads were then split and incubated overnight at 4°C with either 1:100 IgG or α-FLAG primary antibody (Sigma-Aldrich M2 for experiment-1 and ThermoFisher FG4R for experiment-2). After washing to remove unbound primary Ab, pAG-MNase was added to the nuclei and incubated for an hour. Targeted chromatin digestion by pAG-MNase was initiated by adding CaCl_2_ and stopped after 30 mins with stop buffer spiked with 50 ng of *E. coli* DNA. The supernatant with the pAG-MNase digested DNA was then collected and purified using an Epicypher DNA cleanup column. DNA libraries were prepared using the NEB Ultra II protocol kit, with slight modifications as recommended in the Epicypher CUTNRUN kit.

Our pilot experiment (Experiment 1) was set up with U04 strains expressing one of the three alleles (*HF-MRR1^ancestral^*, *HF-MRR1^Y813C^* and *HF-MRR1^L1Q1*^*) and sequenced using paired-end 150-bp reads on the Illumina Nextseq 2000 platform to achieve a sequencing depth of 10M per sample. Based on the pilot study results, the sequencing depth was adjusted to 5-6M per sample for the subsequent experiment (Experiment 2) including two biological replicates of U04 strain expressing *HF-MRR1^Y813C^*, which were sequenced using paired-end 50 bp reads on the Illumina Nextseq 2000 platform.

### CUT&RUN data analysis

Raw read quality was evaluated using FastQC (v0.12.1) prior to read trimming with Cutadapt (v.4.4) for adapter sequences with additional parameters “--nextseq-trim 20-- max-n 0.8--trim-n-m 1”. Reads were mapped to *Clavispora* (*Candida*) *lusitaniae* strain L17 (NCBI accession: ASM367555v2) with Bowtie2 (v2.4.2) using parameters “--local-- no-mixed--no-discordant”. Alignments were sorted coordinate with Samtools (v1.11), filtered for unmapped or multi-mapping reads using sambamba (v0.8.0), and downsampled to 3 million reads per sample to ensure equal sensitivity for peak calling across samples. MarkDuplicates (Picard Tools) was used to identify and remove duplicate reads. Fragment size distributions of individual samples were visualized using deepTools (v3.5.1) command “bamPEFragmentSize”. Peaks were called using the MACS2 (v2.2.7.1) command “callpeak” in narrowpeak mode using IgG IP samples as controls with parameters “-f BAMPE--keep-dup all-g 11999093-q 0.05”. Significant peaks were further filtered to keep only those with 2-fold or greater signal increase relative to control (IgG) samples. The fraction of reads in peaks (FRiP) was calculated for each sample to assess individual quality. The BEDTools (v 2.31.1) command “merge”, with the parameter “-c” for averaging peak signal value, was used to merge peaks with a 2-fold or greater signal from all replicates of experiment-2. BEDTools (v 2.31.1) command “intersect” with the parameter “-a” was used to identify a set of reproducible overlapping peaks between HF-Mrr1^Y813C^ from Experiment 1 and Experiment 2. Since the nuclei isolation step was not controlled in our CUT&RUN experiments (36, 69), it limited our ability to perform differential peak analysis between replicates and across strains expressing different *MRR1* alleles.

### Drug susceptibility assays

Minimum inhibitory concentration (MIC) was determined using a broth microdilution method as previously described (70). Briefly, 2×10^3^ cells were added to a two-fold dilution series of the drug prepared in RPMI-1640 then incubated at 37 °C. The MIC was defined as the minimum drug concentration that abolished visible growth compared to a drug-free control. The MIC_90_ was defined as the minimum drug concentration that led to a 90% or greater decrease in growth relative to a drug-free control. No more than a 2-fold difference was observed between MICs recorded at 24 and 48 h; data from the 24 h timepoint was reported unless otherwise noted. The concentration range used for azoles were FLZ: 64 to 0.0625 µg/ml, VOR: 4 to 0.004 µg/ml, KTZ: 1 to 0.004 µg/ml, ITR: 0.4 to 0.003125 µg/ml and ISA: 1 to 0.04 µg/ml. For the broad-spectrum antifungals, the following concentration ranges were used; Myclobutanil: 32 to 0.0625 µg/ml, Terbinafine: 64 to 0.125 µg/ml, Cycloheximide: 32 to 0.0625 µg/ml, 5-FC: 4 to 0.008 µg/ml, Fluphenazine: 200 to 0.39 µg/ml and Mycophenolic acid: 256 to 0.5 µg/ml.

### MOTIF analysis

Sequences spanning ±100 bp around the peak summits identified from CUT&RUN data were extracted from the L17 genome (NCBI accession: ASM367555v2) using BEDTools v2.30.0 (71). To establish a background control, we used BEDTools random to retrieve randomly selected 200bp sequences from the genome of L17. STREME (43), part of the MEME Suite (streme--verbosity 1--oc streme_results--dna--totallength 4000000--time 14400--minw 6--maxw 20--thresh 0.05--align center--p around_peaks.fasta --n random_sequences.fasta) was employed for motif discovery and enrichment analysis. Motif scanning across *C. lusitaniae* strains (L17, AR0398, ATCC 42720, 79-1, and 76-31) and multiple related *Candida* species (*C. albicans* SC5314 (ASM18296v3), *C. parapsilosis* CDC317 (ASM18276v2), *C. auris* B11205 (ASM1677213v1) or B8441 (GCA_002759435.3), and *C. lusitaniae* ATCC 42720) was conducted using FIMO (72), part of the MEME Suite (fimo --oc fimo_results --verbosity 1 --bgfile --nrdb --thresh 1.0E-3 motif1.meme target_seqs.fasta). *MDR1* and *CDR1* gene IDs and their translational start site coordinates used for the sequence retrieval of the upstream regions are listed in Table S4.

For the phylogenetic gene trees, nucleotide sequences of the respective genes were extracted and aligned using MAFFT v7 (https://mafft.cbrc.jp/alignment/server/index.html) with default parameters. A neighbor-joining tree was then constructed based on the aligned DNA sequences, with 1000 bootstrap replicates to assess phylogenetic relationships. The results were visualized using the ggmotif v0.2.0 R package (73) and FigTree v1.4.4 (http://tree.bio.ed.ac.uk/software/figtree/).

### Statistical analysis and figure design

Ordinary one-way ANOVA and Dunnett’s multiple comparisons testing, with a single pooled variance, were used for statistical evaluation. P values <0.05 were considered significant for all analyses performed and are indicated with asterisks: *p<0.05, **p<0.01, ***p<0.001 and ****p<0.0001. Fig. 6A and 7 were created in BioRender https://BioRender.com/39wn5ht.

## Data availability

The raw sequence reads from CUT&RUN analysis have been deposited into NCBI sequence read archive under Bioproject PRJNA1251050.The data from this study are available within the paper and in the supplemental material, and accessible in OSF through the following link https://osf.io/4zbv8/ and will be made public upon publication.

## Acknowledgements

We thank Owen Wilkins and Noelle Kosarek for CUT&RUN data analysis, Mohammad Qasim, Fred Kolling, Heidi Trask and Jen Spengler for guidance with CUT&RUN pilot experiments and Stacie Stuut for azole structures.

Research reported in this publication was supported by National Institutes of Health (NIH) grant R01 AI127548 to D.A.H. This work was also supported by the Cystic Fibrosis Foundation Research Development Program (CFFRDP) STANTO19R0 for the Translational Research Core. Equipment used was supported by the NIH NIGMS grant to Dartmouth BioMT P20-GM113132. J.E.S. and C.G.P.P. were supported by R01 AI130128. J.E.S. is a CIFAR Fellow in the program Fungal Kingdom: Threats and Opportunities. Sequencing was carried out in the Genomics and Molecular Biology Shared Resource (RRID:SCR_021293) at Dartmouth which is supported by NCI Cancer Center Support Grant 5P30CA023108 and NIH S10 (1S10OD030242) awards. Data analysis was performed by the Dartmouth’s Center for Quantitative Biology through a grant from the National Institute of General Medical Sciences of the NIH award P20GM130454 and on the UC Riverside High Performance Computing Cluster supported by NSF (MRI-2215705, MRI-1429826) and NIH (1S10OD016290-01A1).

---

**Figure S1: Consensus Mrr1-binding DNA motif in the promoter regions of Mrr1 targets.** (A) The positions of the 14-nt (orange hatches) and 9-nt Mrr1-binding motifs (<u>c</u>onsensus <u>M</u>rr1-<u>b</u>inding <u>m</u>otif or cMBM; blue hatches) in the ∼890 bp upstream intergenic regions of *MDR1* in *C. lusitaniae* L17 *and* ATCC 42720 strains. (B) cMBM location in the 1 kb upstream intergenic regions of *CDR1* from *C. lusitaniae* L17 *and* ATCC 42720. (C) cMBM location in the 1 kb upstream intergenic regions of the *CDR1* homologs of *C. parapsilosis* CDC317, *C. auris* B8441*, C. albicans* SC5314 and *C. lusitaniae* ATCC 42720. The phylogenetic tree was constructed using the *CDR1* nucleotide sequences. (B, C) The intergenic region upstream of *CDR1* in ATCC 42720 is 326 bp. Grey arrow indicates the adjacent ORF *CLUG_03114*.

**Figure S2: Comparison of binding profiles of constitutively active and low-activity Mrr1.** (A-C) Comparison of data from Figures 1D-F and Figures 5A-C to highlight the similarities in peak profiles. HF-Mrr1^Y813C^ (in blue) and HF-Mrr1^ancestral^ (in grey) CUT&RUN read coverage plots normalized per 20 bp bin size. Chromosomal positions of regions containing *MDR1, CDR1* and *FLU1* and adjacent genes are represented to scale with boxes and arrows. Peaks from HF-Mrr1-bound DNA recovered by an α-FLAG antibody and for the non-specific binding control recovered via IgG are shown. Signal indicates the average read density in α-FLAG relative to IgG within the peak region.

**Figure S3: Biochemical and phenotypic analysis of HF-tagged Mrr1 variants.** (A) Western blot of whole cell protein lysates of U04 strains expressing N-terminal 6xHis-3xFLAG-tagged Mrr1 (HF-Mrr1) variants. HF-Mrr1 was probed using an α-FLAG antibody. Mean ± SD of HF-Mrr1 band intensities normalized to total protein (n= 4 biological replicates). (B) FLZ MIC of U04 clinical isolate (native allele *MRR1^Y813C^*) and U04 *mrr1*Δ complemented with untagged or *HF-MRR1* was determined by broth microdilution assays. The data shown represent the mean ± SD from three independent experiments. There were no significant differences observed between data from strains with untagged Mrr1 variants and data from strains with their respective HF-tagged counterparts. Strains with constitutive Mrr1 activity are in bold.

**Figure S4: Global binding profiles of constitutively active and low-activity Mrr1.** Circos plot showing global CUT&RUN-determined Mrr1-binding peaks of HF-Mrr1^Y813C^ (in blue), HF-Mrr1^ancestral^ (in grey) and HF-Mrr1^L1Q1*^ (in orange) in the *C. lusitaniae* L17 genome. Mrr1-binding peaks with a signal ≥2-fold compared to their respective IgG backgrounds and up to 1 kb away from the nearest ORF from Experiment 1 (see Supplemental Files 1, 4 and 5) were used. The genomic positions of the 25 differentially expressed genes that constitute the Mrr1-regulon are marked with the L17 gene IDs.

